# A Pilot Randomized Trial of Oral Magnesium Supplementation on Supraventricular Arrhythmias

**DOI:** 10.1101/215319

**Authors:** Pamela L. Lutsey, Lin Y. Chen, Anne Eaton, Melanie Jaeb, Kyle D. Rudser, James D. Neaton, Alvaro Alonso

## Abstract

**Background:** Magnesium is believed to have a physiologic role in cardiac contractility, and evidence from epidemiologic and clinical studies has suggested that low serum concentrations of magnesium may be associated with increased risk of atrial fibrillation (AF).

**Objective:** As part of the planning effort for a large randomized trial to prevent AF with magnesium supplementation, we conducted a 12-week pilot study to assess adherence to oral magnesium supplementation and matching placebo, estimate the effect on circulating magnesium concentrations, and evaluate the feasibility of using an ambulatory monitoring device (ZioPatch) for assessing premature atrial contractions (PACs), a predictor of AF.

**Design:** Double-blind randomized pilot clinical trial comparing supplementation with 400 mg magnesium oxide daily (versus placebo) over 12 weeks of follow-up. The ZioPatch was applied for 14 days at baseline and the end of follow-up. Adherence to the assigned treatment, and changes in PACs, serum magnesium concentration, glucose and blood pressure were assessed.

**Results:** A total of 59 participants, 73% women and average age 62 years, were randomized. 98% of participants completed follow-up. Those assigned to the magnesium supplement took 75% of tablets as compared to 83% for those in the placebo group. Change in magnesium concentrations was significantly greater for those given magnesium supplement compared to placebo (0.07; 95% confidence interval (CI): 0.03, 0.12 mEq/L; p = 0.002). ZioPatch was worn for an average of 13.0 of the requested 14 days at baseline; at the end of follow-up, the average number of days of monitoring was 13.0 days for the magnesium supplement group and 12.7 days for the placebo group. For log PAC burden (episodes per hour), the average change from baseline was −0.05 (95% CI: −0.31, 0.20) for those randomized to magnesium supplement and 0.04 (95% CI: −0.24, 0.31) for those randomized to placebo (p=0.79 for difference). Gastrointestinal problems were reported by 50% of participants in the magnesium supplement group and 7% in the placebo group. Only one person in the magnesium supplement group and none in the placebo group experienced adverse events which led to treatment discontinuation.

**Conclusions:** In this pilot randomized clinic trial, although gastrointestinal side effects to the magnesium supplement were common, adherence, measured by pill counts, was very good and, as a consequence, magnesium concentrations were greater for those randomly assigned to the magnesium supplement compared to placebo. Participant acceptance of the planned monitoring with ZioPatch was also very good. While the difference in the change in PACs was not significant, this pilot study was small, short-term, and did not include participants at high risk of AF. Thus, we could not reliably evaluate the effect of magnesium supplementation on PACs.

**Clinicaltrials.gov registration:** NCT02837328

## Background

Atrial fibrillation (AF) is a common cardiac arrhythmia characterized by irregular atrial electrical activity. In the US, more than 3 million individuals had AF in 2010, and this figure is expected to more than double by 2050.^1–3^ Current AF treatments, including antiarrhythmic drugs and catheter ablation for rhythm restoration, and oral anticoagulation for the prevention of thromboembolism, have suboptimal efficacy and carry significant risks.^4^ Limitations of the available therapeutic approaches highlight the need for primary prevention interventions.^5,6^ As highlighted in a 2009 NHLBI report^5^ and stressed in a more recent Heart Rhythm Society-sponsored whitepaper,^6^ there is an urgent need to identify new and effective strategies for the primary prevention of AF.

Compelling evidence from numerous lines of inquiry suggests that low concentrations of serum magnesium may be causally associated with AF risk. First, magnesium supplementation is recommended as prophylaxis for the prevention of AF in patients undergoing cardiac surgery. A recent Cochrane systematic review and meta-analysis of randomized trials assessing the efficacy of magnesium supplementation for AF prevention in heart surgery reported an odds ratio of 0.55 (95%CI: 0.41, 0.73) for AF or supraventricular arrhythmia comparing the magnesium intervention to control.^7^ Second, indirect evidence from three prospective epidemiologic studies provides some support for such intervention; each reported that individuals in the lowest versus highest quantile of serum magnesium were 35-50% more likely to develop incident AF, after multivariable adjustment.^8–10^ Finally, additional evidence for an effect of magnesium on the risk of arrhythmias is provided by a study of dietary magnesium restriction in which 3 out of 14 women fed a low-magnesium diet developed AF, which resolved quickly after magnesium repletion.^11^

Whether magnesium supplementation could have a role in prevention of AF in the community has not been tested. Were magnesium supplementation shown to prevent AF and be safe over the long-term, it would be an ideal intervention for primary prevention as it is easy to implement and inexpensive.

As part of the planning effort for a large randomized trial to prevent AF with magnesium supplementation, we conducted a double-blind, placebo-controlled randomized clinical trial of oral magnesium supplementation to assess supplement adherence, side effects, effect on serum magnesium concentration, and feasibility of using an ambulatory monitoring device for identification of arrhythmias.

## Methods

### Study Participants

Participants 55 years of age or older were recruited using fliers, the University of Minnesota StudyFinder website, invitations to individuals enrolled in the ResearchMatch research volunteer database, and invitations to University of Minnesota School of Public Health employees. Exclusion criteria included a prior history of heart disease (coronary heart disease, heart failure, AF), stroke, known kidney disease; use of type I or III antiarrhythmic drugs or digoxin; current use of magnesium supplements; any prior history of allergy or intolerance to magnesium; lactose intolerance; prior history of inflammatory bowel disease or any severe gastrointestinal disorder.

Eligible participants attended a baseline visit where measurements were conducted and a Zio^®^ XT Patch (ZioPatch; iRhythm Technologies, Inc., San Francisco, California) heart rhythm monitor was applied by trained staff. After wearing the ZioPatch for 2 weeks, participants were randomized 1:1 to either 400 mg magnesium oxide or placebo using block randomization within two strata of age (younger than 65 and 65 and older). All participants provided written informed consent; the study protocol was approved by the University of Minnesota Institutional Review Board.

Following randomization, participants were mailed the study intervention, which they took for a total of 12 weeks. Ten weeks after beginning the study intervention, participants took part in a follow-up clinic visit, and a second ZioPatch was applied. Participants continued the study intervention until the second ZioPatch was removed (2 weeks after the follow-up clinic visit) (**Figure 1)**.

**Figure 1:**
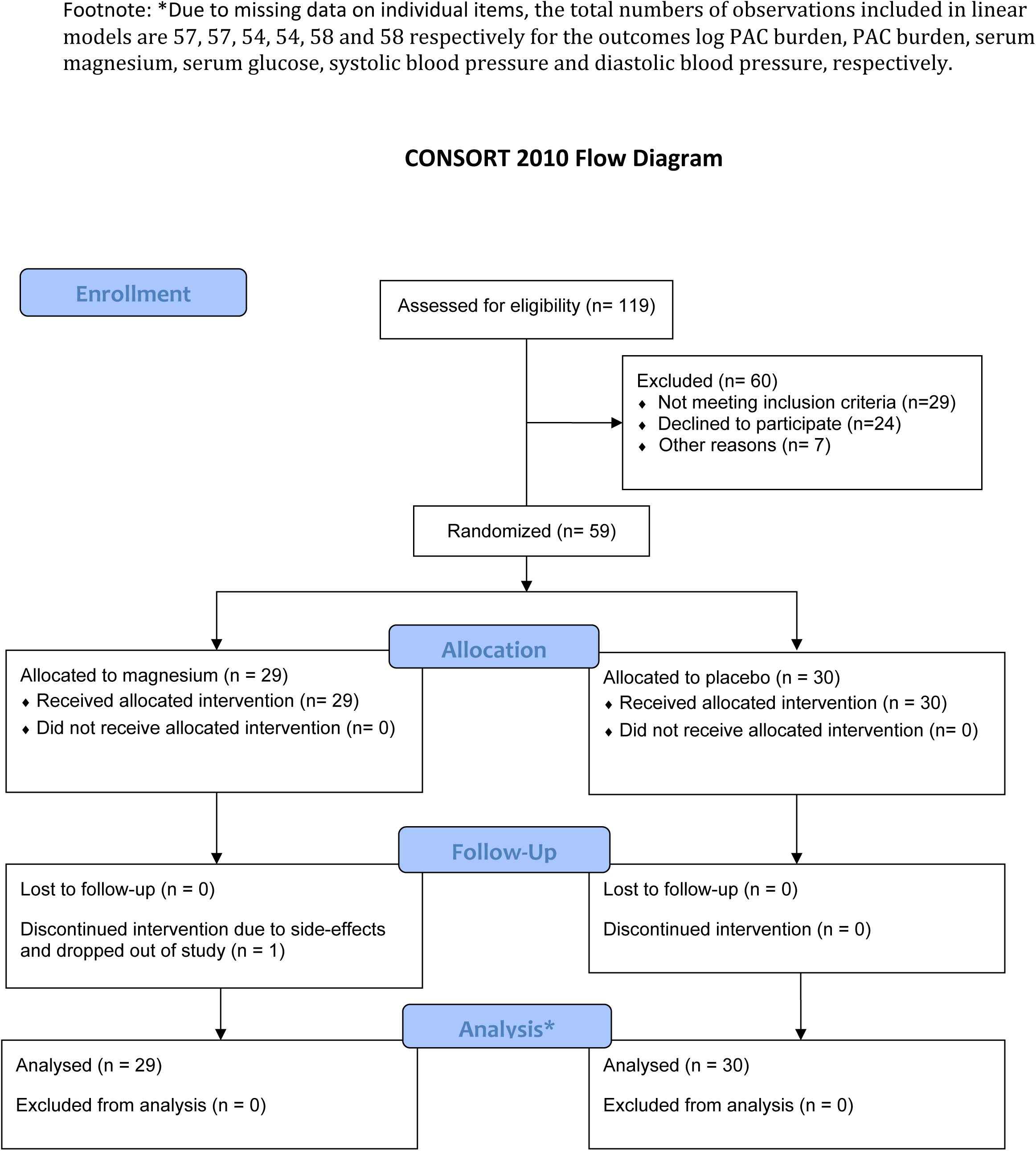

### Study Intervention and Blinding

The University of Minnesota Institute for Therapeutics Discovery and Drug Development produced the active study intervention (400 mg magnesium oxide) and matched placebo (lactose) according to Good Manufacturing Practices. The University of Minnesota Investigational Drug Service managed bottling per the randomization scheme. Study participants and all study staff were blinded to the treatment given.

### Measurements

At the baseline and follow-up clinic visits, participants completed questionnaires and trained study staff conducted physiological measurements (i.e., anthropometry, blood pressure), phlebotomy, and applied the ZioPatch device. Treatment compliance was assessed by a pill count at the follow-up visit. At intervention days 21, 42 and 80 participants were also emailed unique links to online questionnaires, administered via REDCap,^12^ which queried compliance and asked the following open-ended question about adverse effects: “Since starting the study, have you experienced anything out of the ordinary?” Participant blinding was also assessed on day 80, the last day of the study.

The ZioPatch was used to identify premature atrial contractions (PACs). PACs are supraventricular arrhythmias associated with the future risk of AF^13–15^ and are considered an intermediate phenotype of the arrhythmia, reflecting the underlying cardiac substrate that facilitates the development of AF.^16^ Participants were asked to wear the ZioPatches for 2 weeks after each clinic visit. Information obtained from the ZioPatch devices was processed by the ZEUS algorithm, a comprehensive system that analyzes electrocardiographic data received from the device.^17^ We counted as PACs isolated supraventricular ectopic beats, supraventricular ectopic couplet total count, and supraventricular ectopic triplet total count. Total PACs were then divided by number of hours the ZioPatch recorded analyzable data, which yielded PACS per hour.

Participants were asked to fast for 8 hours prior to blood draws. Serum magnesium and glucose were measured using the Roche Cobas 6000 at the University of Minnesota Advanced Research and Diagnostic Laboratory. Blood pressure was measured with the participant sitting, after a 5 minute rest, with a random zero sphygmomanometer (Omron Digital Blood Pressure Monitor HEM-907XL). Three measurements were taken; all three measurements were averaged for use in analyses. Height and weight were measured with participants in light clothing, and shoes removed. Height was measured with a research stadiometer and weight with a scale.

### Statistical Analysis

The goal of the pilot study was to assess adherence to the Mg supplement, and feasibility of using the ZioPatch and collect preliminary data on PACs, a predictor of AF. Sample size was determined to detect a difference in the change in PACs (follow-up minus baseline) between treatment groups of 0.79 standard deviation units with 80% power and 5% type I error (2-sided), assuming 5 participants would not complete follow-up.

All analyses were intent-to-treat. Descriptive statistics are provided according to treatment assignment for baseline characteristics, adherence, Mg concentrations and other outcomes. Linear regression was used to evaluate whether change in outcomes differed according to treatment assignment, adjusting for the randomization stratification factor (age ≥65 vs <65) and baseline value of the outcome with robust variance estimates for confidence intervals and P-values. Post-hoc sensitivity analyses further adjusted for sex. As PAC burden is highly skewed, we pre-specified using log PAC burden for analysis and reported the ratio of geometric means. Pre-specified subgroup analyses were also performed, stratified by baseline magnesium concentration (< vs. ≥ median). A 2-sided p-value of <0.05 was used to indicate statistical significance. Analyses were conducted using R^18^ version 3.4.0 (R Foundation, Vienna, Austria).

## Results

### Study Participants

Between March and June 2017, 59 participants were randomized, 29 to Mg supplement and 30 to matching placebo. Participant characteristics were generally similar by treatment group, with the notable exception of sex; 86.2% of participants in the treatment group were women while in the placebo group 60.0% were female (**Table 1**). The mean age of participants was 61.5 ± 5.2 years. Baseline serum magnesium was 1.74 ± 0.11 mEq/L in participants assigned magnesium supplements and 1.71 ± 0.10 in those assigned placebo; 6.9% had magnesium concentrations below the threshold for clinical deficiency (<1.5 mEq/L) while 37.9% had concentrations below the threshold for subclinical deficiency (<1.7 mEq/L).

**Table 1:** Baseline participant characteristics^*^ overall and stratified by intervention status

|  | Overall | Magnesium<br>(400 mg daily) | Placebo |
| --- | --- | --- | --- |
| *N* | 59 | 29 | 30 |
| <u>*Demographics*</u> |  |  |  |
| Age, years | 61.5 ± 5.2 | 61.3 ± 5.3 | 61.6 ± 5.2 |
| Age category |  |  |  |
| ≥ 65 years | 14 (23.7) | 6 (20.7) | 8 (26.7) |
| < 65 years | 45 (76.3) | 23 (79.3) | 22 (73.3) |
| Sex |  |  |  |
| Female | 43 (72.9) | 25 (86.2) | 18 (60.0) |
| Male | 16 (27.1) | 4 (13.8) | 12 (40) |
| Race |  |  |  |
| White | 56 (94.9) | 27 (93.1) | 29 (96.7) |
| Nonwhite | 3 (5.1) | 2 (6.9) | 1 (3.3) |
| Educational attainment |  |  |  |
| High school graduate or GED | 1 (1.7) | 0 (0) | 1 (3.3) |
| Some college | 10 (16.9) | 6 (20.7) | 4 (13.3) |
| College graduate | 26 (44.1) | 10 (34.5) | 16 (53.3) |
| Graduate school or<br>professional school | 22 (37.3) | 13 (44.8) | 9 (30) |
| <u>*Physiologic characteristics*</u> |  |  |  |
| BMI, kg/m<sup>2</sup> | 27.9 ± 4.64 | 27.7 ± 4.9 | 28.0 ± 4.5 |
| Serum magnesium, mEq/L | 1.72 ± 0.11 | 1.74 ± 0.12 | 1.71 ± 0.10 |
| Systolic blood pressure, mmHg | 119 ± 16 | 119 ± 14 | 119 ± 18 |
| Diastolic blood pressure, mmHg | 71 ± 9 | 72 ± 9 | 71 ± 10 |
| Antihypertensive medication | 14 (24) | 9 (31) | 5 (17) |
| Serum glucose, mg/dL | 98.9 ± 29.9 | 94.2 ± 10.6 | 103.2 ± 40.2 |
| Sensitivity analysis** | 95.2 ± 11.1 | 94.2 ± 10.6 | 96.2 ± 11.6 |
| Glucose lowering medication | 2 (3.4) | 0 (0) | 2 (6.7) |
| PAC burden, episodes/hour | 13.9 ± 56 | 8.1 ± 14 | 19.5 ± 78 |
| [25<sup>th</sup>, 75<sup>th</sup> percentiles] | [1.07, 6.01] | [1.18, 6.99] | [1.03, 3.69] |
| Log PAC burden, log(episodes/hour) | 1.04 ± 1.46 | 1.14 ± 1.40 | 0.95 ± 1.53 |

\*mean ± SD or n (%)
\*\*Omission of one participant with a baseline glucose value of 307 mg/dL

Log PAC burden (episodes per hour) at baseline was 1.14 ± 1.40 in the treatment group and 0.95 ± 1.53 in the placebo group. At baseline, average ZioPatch analyzable time in the intervention and placebo groups were 13.1 ± 1.7 and 12.9 ± 2.6 days, respectively, with 93.1% assigned to magnesium and 90.0% assigned to placebo wearing ≥12 days.

### Follow-up

A total of 2 participants, both in the intervention group, were missing ZioPatch information at follow-up; 1 participant dropped out of the study and 1 due to a device malfunction (Figure 1).

### Adherence and Mg Concentrations

Based on pill count, participants in the magnesium group took 75.1% ± 17.8% of tablets, whereas those in the placebo group took 83.4% ± 5.9%. Self-reported information about the percent missing pills, and reasons for missing pills, is provided in **Table 2**.

**Table 2:** Self-reported compliance.

| Compliance | % reporting missing pills in: |  |  | Reason for missing pills, N |  |  |  |
| --- | --- | --- | --- | --- | --- | --- | --- |
|  | Last 3 days | Last week | Last 2 weeks | Forgot | Too busy | Makes me sick | Other |
| Intervention Day 21 |  |  |  |  |  |  |  |
| Mg | 8% | 12% | 20% | 2 | 0 | 2 | 1 |
| Placebo | 0% | 7% | 24% | 7 | 1 | 0 | 0 |
| Intervention Day 42 |  |  |  |  |  |  |  |
| Mg | 22% | 33% | 39% | 5 | 0 | 5 | 1 |
| Placebo | 15% | 22% | 31% | 6 | 2 | 0 | 2 |
| Intervention Day 80 |  |  |  |  |  |  |  |
| Mg | 14% | 27% | 40% | 5 | 0 | 1 | 0 |
| Placebo | 12% | 23% | 28% | 6 | 1 | 0 | 1 |

Over the 12 week follow-up period those assigned magnesium supplementation had a significant increase in serum magnesium concentration as compared to placebo (0.07 mEq/L; 95% CI: 0.03, 0.12; p=0.002) (**Table 3)**. In subgroup analyses change in magnesium concentration did not vary by baseline magnesium concentration (**Table 4;** interaction 0.24). Specifically, among participants who at baseline were below the median serum magnesium concentration (i.e. 1.74 mEq/L), the effect of magnesium versus placebo on change in serum magnesium concentration was 0.05 (95% CI: 0.00, 0.10) whereas among those at or above the median at baseline the effect was 0.12 (95% CI: 0.04, 0.20).

**Table 3:** Change in PACs, and secondary endpoints (i.e. SBP, DBP and serum glucose, Mg) according to treatment group

|  | Magnesium<br>(400 mg daily)<br>Mean (SD) | Placebo<br>Mean (SD) | Intervention effect<br>coefficient* (95% CI) | p-value |
| --- | --- | --- | --- | --- |
| Primary outcome (episodes/hr) |  |  |  |  |
| Log PAC burden |  |  | 0.96 (0.70, 1.31)** | 0.79 |
| Baseline | 1.14 (1.4) | 0.95 (1.53) |  |  |
| Follow-up<sup>†</sup> | 1.06 (1.46) | 0.98 (1.58) |  |  |
| Change | -0.05 (0.65) | 0.04 (0.74) |  |  |
| PAC burden |  |  | 0.49 (-2.5, 3.48) |  |
| Baseline | 8.1 ± 14 | 19.5 ± 78 |  |  |
| Follow-up<sup>†</sup> | 7.7 ± 12 | 14.2 ± 47 |  |  |
| Change | -0.5 ± 7 | -5.4 ± 32 |  |  |
| Secondary outcomes |  |  |  |  |
| Serum magnesium, mEq/L |  |  | 0.07 (0.03, 0.12) | 0.002 |
| Baseline | 1.74 (0.12) | 1.71 (0.1) |  |  |
| Follow-up<sup>‡</sup> | 1.8 (0.13) | 1.71 (0.11) |  |  |
| Change | 0.07 (0.09) | 0 (0.1) |  |  |
| Serum glucose, mg/dL |  |  |  |  |
| Baseline | 94.2 (10.6) | 103.2 (40.2) | 2.4 (-3.0, 7.7) | 0.39 |
| Follow-up<sup>‡</sup> | 96.3 (12.2) | 96.2 (13.7) |  |  |
| Change | 1.8 (7.5) | -7.1 (32.8) |  |  |
| Serum glucose<sup>¥</sup>, mg/dL |  |  |  |  |
| Baseline | 94.2 (10.6) | 96.2 (11.6) | 2.8 (-0.9, 6.4) | 0.14 |
| Follow-up<sup>‡</sup> | 96.3 (12.2) | 95 (12.4) |  |  |
| Change | 1.75 (7.5) | -1.21 (6.7) |  |  |
| Systolic blood pressure, mmHg |  |  | 2.9 (-1.8, 7.2) | 0.18 |
| Baseline | 119 (14) | 119 (18) |  |  |
| Follow-up<sup>‡</sup> | 118 (14) | 115 (14) |  |  |
| Change | -1 (10) | -4 (10) |  |  |
| Diastolic blood pressure, mmHg |  |  | -0.5 (-3.5, 2.5) | 0.74 |
| Baseline | 71.8 (8.7) | 70.6 (10.3) |  |  |
| Follow-up<sup>‡</sup> | 71.0 (8.8) | 70.8 (8.7) |  |  |
| Change | -0.5 (7.1) | 0.2 (6.1) |  |  |

\*Adjusted for age ( $\geq 65$ or $< 65$ ), and baseline concentration (e.g. when change in glucose is the outcome, models were adjusted for baseline glucose). The numbers of observations included in linear models are 57, 57, 54, 54, 53, 58 and 58 respectively for the outcomes log PAC burden, PAC burden, serum magnesium, serum glucose, serum glucose excluding outlier, systolic blood pressure and diastolic blood pressure, respectively.
\*\* Presented as a ratio of geometric means, i.e. $\exp(\text{coefficient})$
<sup>†</sup>ZioPatch was worn for a 2-week period, from the follow-up clinic visit (intervention week 10) through the end of the study (intervention week 12)
<sup>‡</sup>Follow-up information obtained at clinic visit (intervention week 10)
<sup>¥</sup>Outlier removed.

**Table 4:**
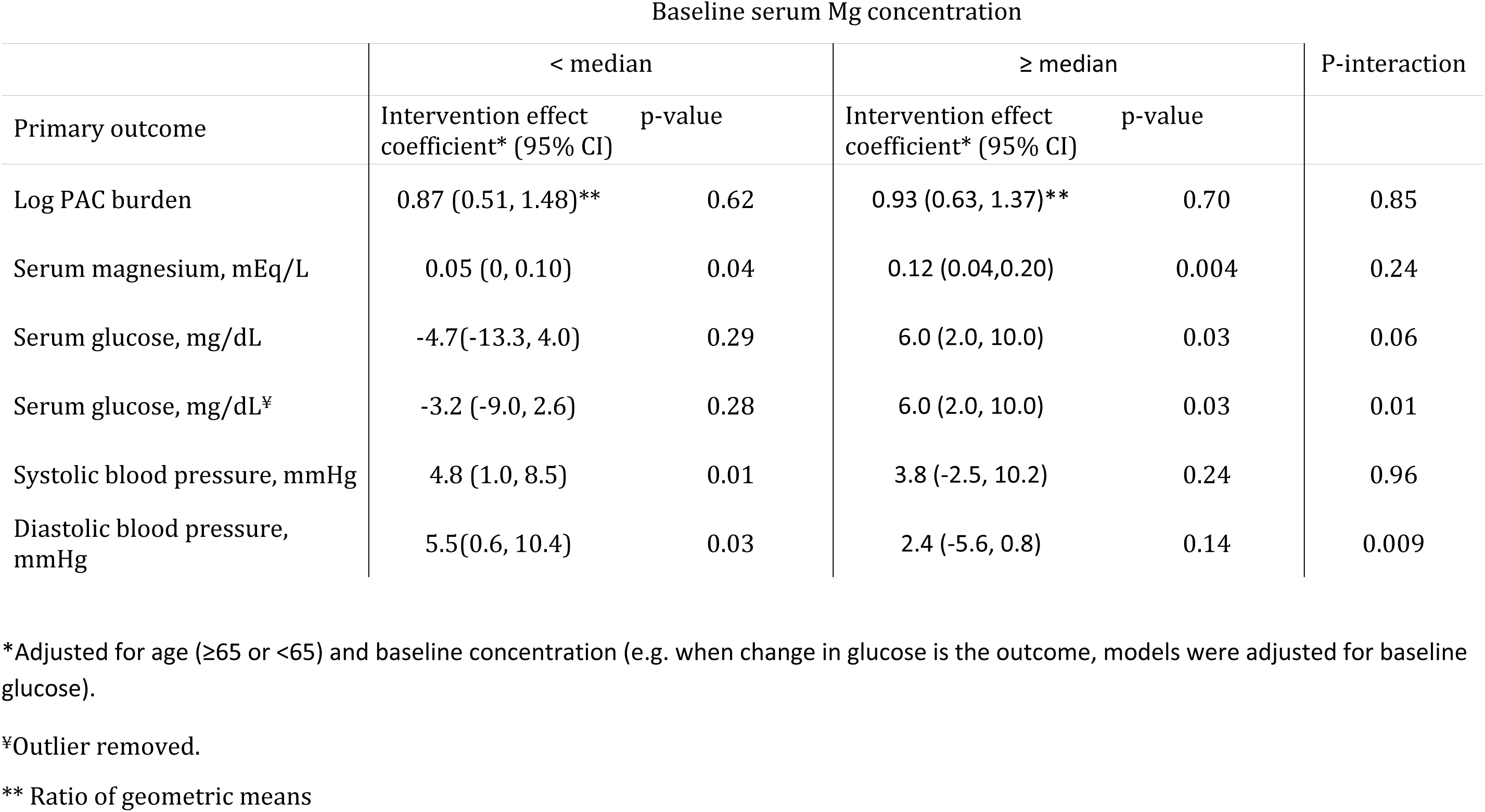
Change in PACs, and secondary endpoints (i.e. SBP, DBP and serum glucose, Mg) according to treatment group, stratified by baseline serum Mg concentration.

| Baseline serum Mg concentration |  |  |  |  |  |
| --- | --- | --- | --- | --- | --- |
|  | < median |  | ≥ median |  | P-interaction |
| Primary outcome | Intervention effect coefficient* (95% CI) | p-value | Intervention effect coefficient* (95% CI) | p-value |  |
| Log PAC burden | 0.87 (0.51, 1.48)** | 0.62 | 0.93 (0.63, 1.37)** | 0.70 | 0.85 |
| Serum magnesium, mEq/L | 0.05 (0, 0.10) | 0.04 | 0.12 (0.04,0.20) | 0.004 | 0.24 |
| Serum glucose, mg/dL | -4.7(-13.3, 4.0) | 0.29 | 6.0 (2.0, 10.0) | 0.03 | 0.06 |
| Serum glucose, mg/dL<sup>‡</sup> | -3.2 (-9.0, 2.6) | 0.28 | 6.0 (2.0, 10.0) | 0.03 | 0.01 |
| Systolic blood pressure, mmHg | 4.8 (1.0, 8.5) | 0.01 | 3.8 (-2.5, 10.2) | 0.24 | 0.96 |
| Diastolic blood pressure, mmHg | 5.5(0.6, 10.4) | 0.03 | 2.4 (-5.6, 0.8) | 0.14 | 0.009 |

\*Adjusted for age (≥65 or <65) and baseline concentration (e.g. when change in glucose is the outcome, models were adjusted for baseline glucose).
<sup>‡</sup>Outlier removed.
\*\* Ratio of geometric means

### Effect of magnesium supplementation on trial outcomes

**Table 3** presents study outcome values at baseline and follow-up, as well as age- and baseline value-adjusted differences in change according to intervention assignment. Spaghetti plots depicting individual change over the intervention period are provided in **Figure 2**. At follow-up, average wear times were similar to baseline, with 13.0 ± 1.8 days for the intervention group, 12.7 ± 2.3 days for the placebo group, and 92.6% assigned to magnesium and 73.3% assigned to placebo wearing ≥12 days. For the primary outcome, log PAC burden (episodes per hour), change over the intervention period was −0.05 (95% confidence interval (CI): −0.31, 0.20) for those randomized to magnesium supplement and 0.04 (95% CI: −0.24, 0.31) for those randomized to placebo. In multivariable-adjusted models there was no evidence of an intervention effect; the ratio of geometric means was 0.96 (95% CI: 0.70, 1.31), p-value = 0.79. Similarly, in subgroup analyses the effect did not differ according to baseline magnesium concentration above versus below the median (**Table 4**; p-interaction = 0.85).

**Figure 2:**
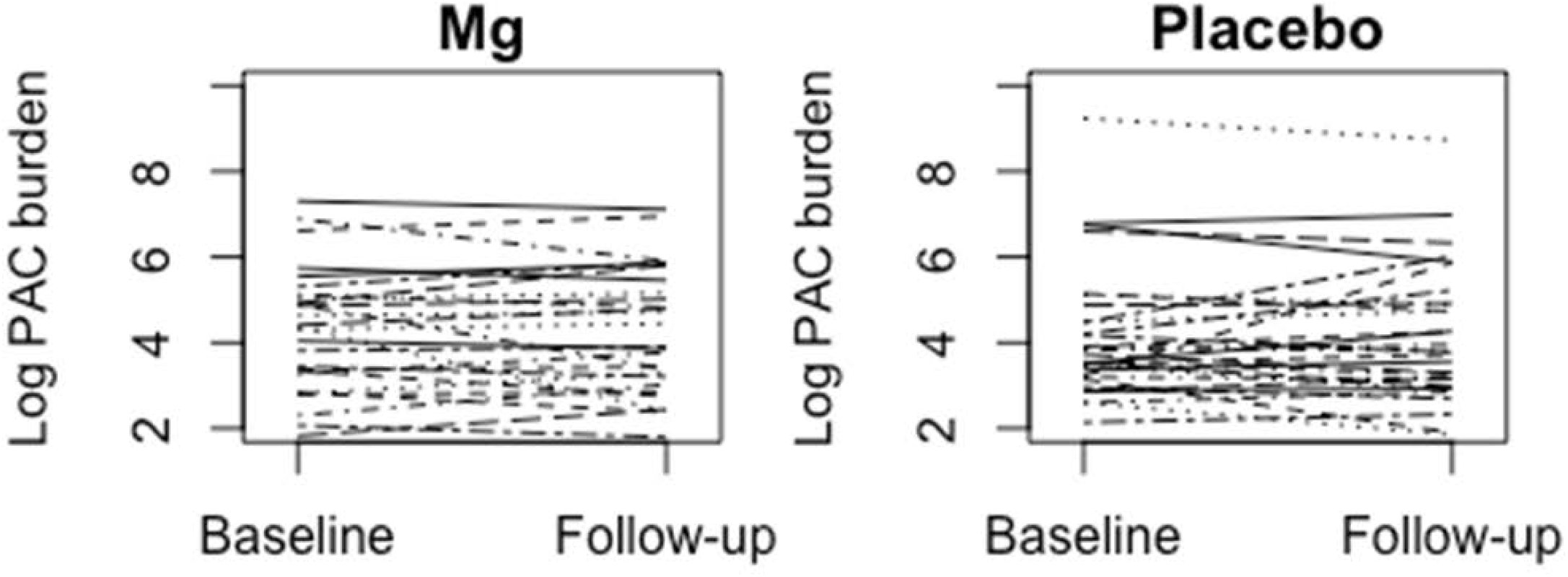
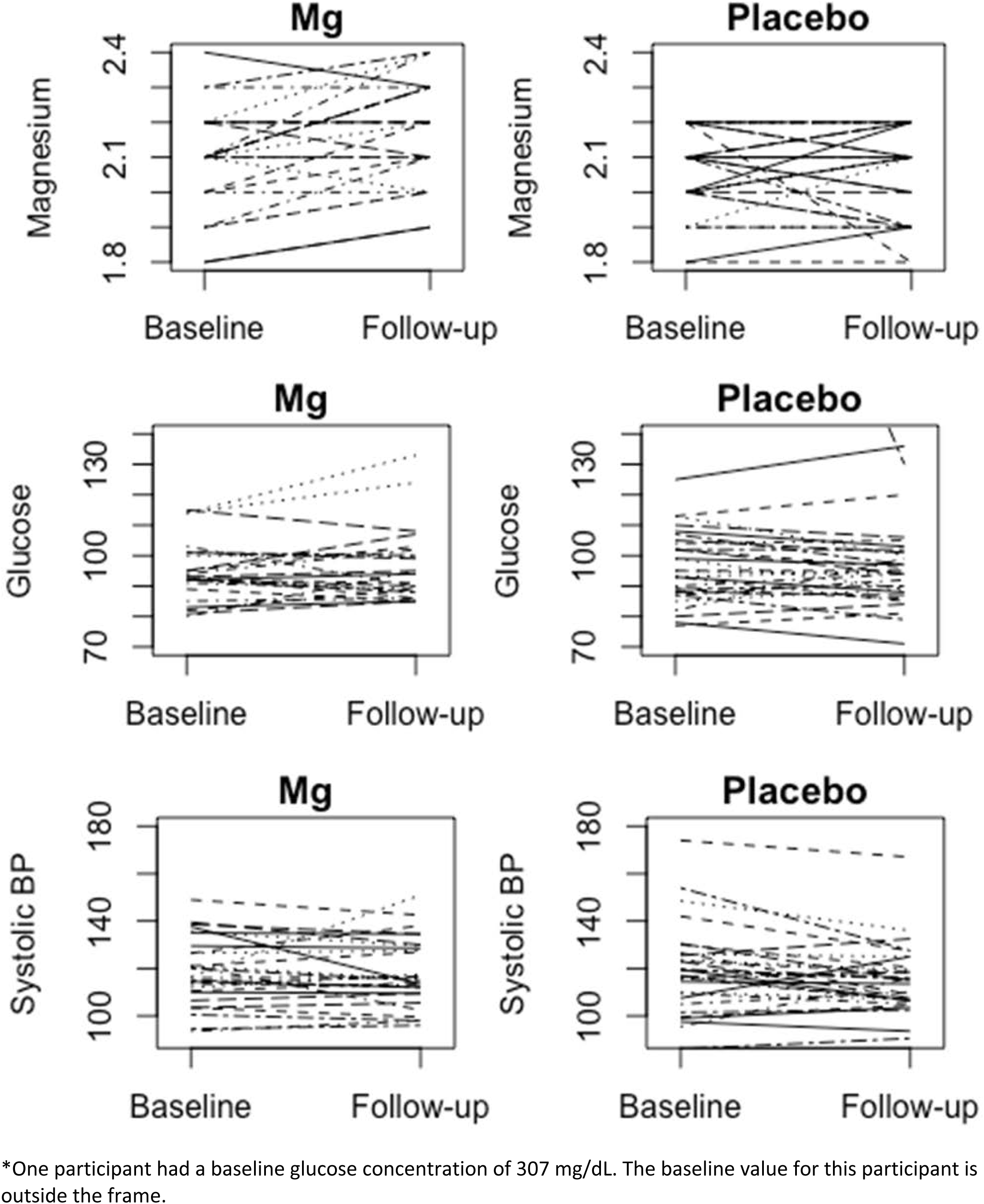
Spaghetti plots for change in log PAC burden, change in serum Mg, serum glucose^*^ and SBP.

Magnesium supplementation was not significantly associated with change in serum glucose [2.4 (95% CI: −3.0, 7.7) mg/dl; p = 0.39]. The lack of association remained in sensitivity analyses excluding one participant with extremely high baseline glucose [2.8 (95% CI: −0.9, 6.4) mg/dl] and one participant who reported changing his/her diabetes medication status during follow-up [2.0 (95%: −3.6, 7.5) mg/dl]. The intervention was also not significantly associated with change in systolic or diastolic blood pressure overall [2.9 (95%: −1.4, 7.2) mmHg and −0.5 (95%: −3.5, 2.5) mmHg, respectively], or after excluding two participants who changed their blood pressure medication status between the baseline and follow-up visits (data not shown).

In post-hoc analyses where we additionally adjusted for sex, results were similar (data not shown). Also, no meaningful patterns emerged in additional subgroup analyses by age category and sex (data not shown).

### Safety and tolerability of the intervention

When asked an open-ended question about adverse events, the most commonly reported responses related to gastrointestinal (GI) symptoms. Of the intervention group, 32% commented on GI changes at intervention day 21, 30% at day 42, and 33% at day 80. In the placebo group 7% commented on GI changes at day 21, 4% at day 42, and 0% at day 80. When considering unique individuals, 50% assigned to magnesium and 7% assigned to placebo commented on GI changes at any point in the study. Specific GI comments, by study participation day, are provided in **Supplemental Table 1**.

One person in the intervention group experienced side effects which led the participant to discontinue blinded study treatment.

At the end of the study, when asked what group they were assigned to, among those assigned to the active treatment, 15% guessed magnesium supplements, 14.3% placebo, and 35.7% reported not knowing (15 did not respond). Of those assigned to placebo, 4.3% guessed magnesium supplements, 26.1% placebo, and 69.6% reported not knowing (7 did not respond).

## Discussion

In this pilot trial of 59 relatively healthy adults aged 55 and older, supplementation with 400 mg magnesium daily over 12 weeks was safe and well-tolerated, and led to a change of 0.07 mEq/L in serum magnesium, which is substantial enough in magnitude that in a larger sample size may translate to health outcomes. The intervention was not associated with change in PACs, but estimates of association had wide confidence intervals and the study was not powered to identify important differences. Likewise, there was no association between supplemental magnesium and change in glucose, systolic blood pressure, or diastolic blood pressure.

The mechanisms through which magnesium supplementation could reduce the risk of supraventricular arrhythmias and AF are not fully understood. However, magnesium is known to play a direct role in cardiac contractility.^19^ Small studies in healthy individuals and in patients with cardiac disease have found that intravenous magnesium administration prolongs sinoatrial, intra-atrial, and atrioventricular node conduction, and the atrial refractory period, which in turn may contribute to prevent onset of AF.^20–22^ Also, randomization to 148 mg oral magnesium (and 296 mg potassium) intake (vs. placebo) had antiarrhythmic effects among 232 patients with frequent ventricular arrhythmias.^23^

Blood pressure and diabetes are also established risk factors for AF,^24,25^ through which magnesium may lower AF risk. In the present pilot trial changes in blood pressure and serum glucose did not differ significantly between those given magnesium supplementation and those given placebo. This is in contrast to the existing literature; however, our study was small and confidence intervals around the treatment differences were wide. Meta-analyses of RCTs have consistently demonstrated that magnesium supplementation lowers blood pressure in a dose-dependent manner.^26–28^ In the most recent meta-analysis, a median dose of 368 mg/d for a median duration of 3 months significantly reduced systolic BP by 2.0 mm Hg (95% CI: 0.4, 3.6) and diastolic BP by 1.8 mm Hg (95% CI: 0.7, 2.8).^28^ Based partly on this evidence, in November 2016 a petition was filed with the Food and Drug Administration (FDA) for a qualified health claim for magnesium and reduced risk of high blood pressure (FDA-2016-Q-3770). A comparable meta-analysis of RCTs, including a total of 370 patients with type 2 diabetes (median intervention duration of 12 weeks, median oral magnesium dose of 360 mg/day), found that magnesium supplementation reduced levels of fasting blood glucose (−10.1 mg/dl, 95%CI – 19.8, −0.2).^29^ These meta-analyses suggest that magnesium is causally related to hypertension and abnormal glucose homeostasis. However, their interpretation is complicated by the fact that the individual studies included in the meta-analyses were highly heterogeneous in terms of magnesium formulation and dosage, and participant characteristics.

In terms of serum magnesium, the intervention of 400 mg magnesium oxide daily was associated with a serum increase of 0.07 mEq/L. This finding is concordant with results from a meta-analysis of the effect of magnesium supplementation dosage on serum magnesium response. In the meta-analysis the median dose was 360 magnesium/day, intervention length 12 weeks and response 0.08 mEq/L.^30^ In the meta-analysis there was evidence of an inverse relationship between baseline magnesium concentration and responsiveness to supplementation. A similar phenomenon was not observed in the present trial, however in our sample baseline magnesium concentrations were quite high and power was exceedingly low for subgroup comparisons.

Results from this study provide additional evidence about the compliance with magnesium supplementation at the dosage of 400 mg magnesium daily, and also safety and tolerability. Among the participants randomized to magnesium only 1 of 29 participants (3.5%) ceased the intervention due to side-effects. Compliance in this study was good, at 75% in the intervention group and 83% in the placebo group, according to pill counts. The low drop-out rate and high compliance provides support for the tolerability of this dosage. However, the fact that 50% in the intervention group commented on GI changes at some point in follow-up should not be dismissed. Unfortunately, given the way side-effects were assessed, it is not possible to quantify the severity of GI complaints. Notably, several individuals only commented about GI changes the first few days after taking the study treatment.

The primary limitation of this study is the small size. An additional consideration is that baseline serum concentrations of the trial participants were quite high; it is unclear how serum magnesium would have changed in a context of low magnesium, and how that may translate to change in other physiologic outcomes. Lastly, we assessed tolerability with a simple open-ended question, not a checklist of specific signs and symptoms graded for severity according to a standard toxicity table.

In sum, this small pilot double-blinded randomized controlled trial of supplementation with 400 mg magnesium daily provides evidence to support the safety and tolerability of this intervention, and also for adherence to the ZioPatch heart rhythm monitoring device. Despite our study population being largely magnesium replete, a change in serum magnesium was observed. Magnesium supplementation was not associated with change in PACs, glucose or blood pressure, however this small pilot study was short-term and not powered to identify small to moderate clinically relevant differences. Results of this pilot study will guide the design of a large trial to evaluate the effect of supplemental magnesium on arrhythmias.

